# Moderate prenatal alcohol exposure increases total length of L1 axons in E15.5 mice

**DOI:** 10.1101/2020.07.15.204842

**Authors:** Shannon Kiss, Karen Herrera, Abigail McElroy, Kelsey Blake, Avery Sicher, Emily Crocker, Christa Jacob, McKayla Lefkove, Myla Cramer, Allysen Henriksen, Josef Novacek, Jenna Severa, Justin Siberski, Emily Thomas, Carlita B. Favero

**Affiliations:** Ursinus College, Department of Biology and Neuroscience Program, Collegeville, PA

**Author notes:** Correspondence to: Carlita B. Favero.

## Abstract

Public health campaigns broadcast the link between heavy alcohol consumption during pregnancy and physical, cognitive, and behavioral birth defects; however, they appear less effective in deterring moderate consumption prevalent in women who are pregnant or of childbearing age. The incidence of mild Fetal Alcohol Spectrum Disorders (FASD) is likely underestimated because the affected individuals lack physical signs such as retarded growth and facial dysmorphology and cognitive/behavioral deficits are not commonly detected until late childhood. Sensory information processing is distorted in FASD, but alcohol effects on the development of axons that mediate these functions are not widely investigated. We hypothesize that alcohol exposure alters axon growth and guidance contributing to the aberrant connectivity that is a hallmark of FASD. To test this, we administered alcohol to pregnant dams from embryonic day (E) 7.5 to 14.5, during the time that axons which form the major forebrain tracts are growing. We found that moderate alcohol exposure had no effect on body weight of E15.5 embryos, but significantly increased the length of L1+ axons. Our findings support our hypothesis. Future studies will investigate cellular, molecular, and functional mechanisms that underlie these effects.

## Introduction

Fetal alcohol spectrum disorder (FASD) is characterized by cognitive defects, distinct facial morphology (on the severe end of the spectrum), and growth deficiencies due to prenatal alcohol exposure (Sokol et al, 2003). It has been shown that alcohol use disorders and comorbid mental disorders have been linked to disruptive volumetric changes and deterioration to the thalamus (De Bellis et al, 2005; Harding et al, 2005).

Current literature indicates both structural and functional differences as a result of prenatal alcohol exposure. Studies have indicated a reduction of thalamic volume in children with FASD (Lebel et al, 2008) as well as reductions in the cortex, hypothalamus and thalamus connectivity in rat models of FASD (Rodriguez et al, 2016).

Correct migration of thalamocortical axons to their laminar position is a complex, multistep process that relies on guidance cues to avoid migration into the hypothalamus (Garel and Lopez-Bendito 2014). Studies have shown that prenatal alcohol exposure disrupts this process, leading to a disorganized cortex; this disorganization results in altered communication between brain regions, specifically the thalamocortical connections and crossed corticothalamic connections, contributing to the observed deficiencies due to prenatal alcohol exposure (Granato et al, 1995). The current study operated within the paradigm of moderate prenatal alcohol exposure, equivalent to one glass of wine a night within the first trimester, similar to a paradigm used in Swiss Webster Mice (Chi et al, 2015). It was expected that alcohol-exposed mice brains at E15.5 would demonstrate a reduction in the length of L1+ axons, a marker for both thalamocortical and corticothalamic axonal migration. Differences in embryonic weight due to sex and treatment were also investigated.

## Results

### Gender and/or moderate prenatal alcohol exposure do not affect E15.5 mice weight

In order to explore the effects of prenatal alcohol exposure on thalamocortical axon (TCA) guidance and formation, we immunostained axons with LI-NCAM. There was no effect of sex (*p* = 0.46) or treatment (*p* = 0.17) on embryonic body weight nor was there an interaction between sex and treatment (*p* = 0.895, Two-way ANOVA; Fig 1b). Our results do not support our hypothesis that sex and treatment changes body weight at E15.5.

**Figure 1.**
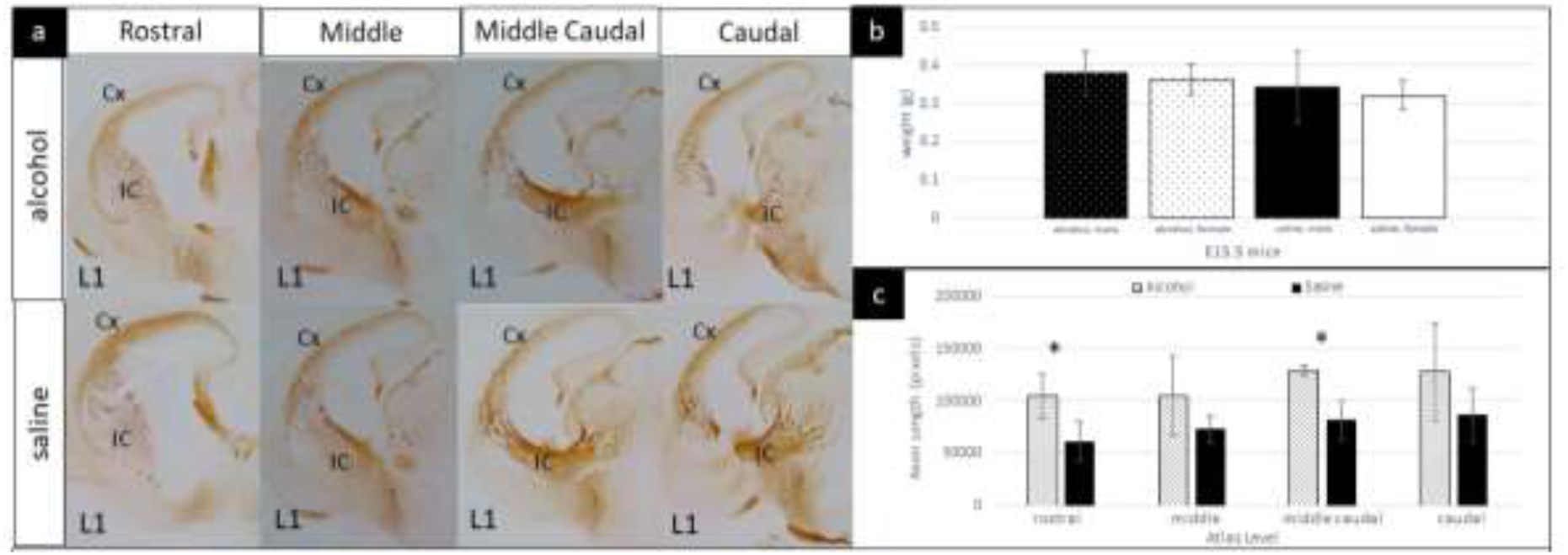
Moderate prenatal alcohol exposure increases total length of L1+ axons, but has no impact on body weight in E15.5 mice. (a) Representative pictures of E15.5 coronal brain sections of alcohol- and saline-exposed groups. Atlas levels are in reference to an E16 mouse brain atlas (Schambra, 2008) in coronal sections. (b) There is no effect of sex (*p* = 0.46) or treatment (*p* = 0.17) on embryonic body weight nor is there an interaction between sex or treatment (*p* = 0.895, Two-way ANOVA; alcohol males: n = 7, 0.38 ± 0.06g, alcohol females: n = 5, 0.36 ± 0.08g, saline males: n = 10, 0.34 ± 0.095g, saline females: n = 8, 0.32 ± 0.04g). (c) Rostral and middle caudal sections showed a significant increase in total length of L1+ axons in the alcohol group compared to the saline group (rostral, alcohol group: n = 3, 102773.83 ± 20703.19, saline group: n = 4, 61342.12 ± 19147.01, *p* = 0.04; middle caudal, alcohol group: n = 4, 129558.24 ± 5358.48, saline group: n = 4, 81814.62 ± 18516.39, *p* = 0.01. There is no interaction between treatment and atlas level (*p* = 0.90, two-way ANOVA). Mean ± S.D. Error bars represent S.D. Statistical analysis was performed with t-test and with a two-way analysis of variance (ANOVA). A post-hoc t-test followed two-way ANOVA for axon length data.

### E15.5 brains show increase in L1+ axon length in response to moderate prenatal alcohol exposure

Utilizing an atlas of an E16 mouse brain (Schambra, 2008) in coronal sections, we refer to each level in relative rostral to caudal directions of the embryonic brain. After visually analyzing L1+ axons and finding no visibly distinct differences across the atlas levels examined (Fig. 1a), we quantified the axon lengths using the AxonTracer plugin for ImageJ (Patel et al., 2018). Among the 4 atlas levels there were no significant differences (*p* = 0.35), between treatments there was a significant difference (*p* = 0.0145), and there was no interaction (*p* = 0.90, Two-way ANOVA). In the middle area of the embryonic brains there was no significant difference in axon length (*p* = 0.26, Two-way ANOVA followed by post-hoc T-test) between the alcohol group (n = 3, 99,052.17 ± 38,276.89) and control group (n = 4, 73,467.62 ± 13,347.25, Fig. 1c). The most caudal area also showed no significant difference in axon length (*p* = 0.14) between the alcohol group (n = 8, 127,992.56 ± 47,209.57) when compared to the control (n = 4, 86,876.12 ± 26,180.88, Fig. 1c). However, in the rostral area of each brain there was a significant increase in axon length in pixels (*p* = 0.04) in the alcohol group (n = 3, 102,773.83 ± 20,703.19) compared to the control group (n = 4, 61,342.12 ± 19,147.01, Fig. 1c). The caudal area of the brains also showed a significant increase in axon length (*p* = 0.01) in embryos exposed to alcohol (n = 8, 129,558.24 ± 5,358.48) compared to those with the saline treatment (n = 4, 81,814.62 ± 18,516.39, Fig. 1c). Our results do not support our hypothesis that axon length would be decreased due to moderate prenatal alcohol exposure but instead show increased axon length.

## Discussion

One important caveat in our interpretation of data has to do with the fact that pregnant dams were subjected to isoflurane anesthesia to minimize distress due to daily injections. It is possible that the changes we observed represent an interaction between anesthesia and maternal alcohol exposure. Our findings are in line with those of Godin et al. that showed a single high dose alcohol exposure on gestational day 7 resulted in a thickened internal capsule with aberrant projections in offspring (Godin et al., 2010). We are unaware of any study investigating axon length in embryos *in vivo*, so we cannot determine if our findings are within range. While previous studies investigating the effect of prenatal alcohol exposure have demonstrated decreased birth weights for exposed embryos, it is likely that there was no observable weight difference in our study due to the moderate level of alcohol exposure compared to the higher level other studies have utilized (Petrelli et al, 2018).

It is well known that alcohol exposure prevents growth cone collapse (Lindsley et al, 2011) and thus influences axonal projection by causing a decrease in cortical growth cone responsiveness (Sepulveda et al, 2011). It is possible that the L1+ axons within our study display a lack of guidance cue responsiveness, resulting in the increased axonal length observed. This is especially pertinent, as it has been hypothesized that organization of TCAs may first begin within the internal capsule, long before the projections reach the cortex (Grove, 2005). Future studies would benefit from investigation into the process of the TCA positioning and organization within the cortex as a result of changes we’ve observed due to moderate prenatal alcohol exposure.

## Methods

### Mice and Injections

Timed pregnant Swiss Webster mice, which were E2.5 upon arrival were purchased from Charles River Laboratories. Mice were single housed and maintained on a 12-hour reverse light-dark cycle. All procedures were approved by the Ursinus College Institutional Animal Care and Use Committee and are in accordance with the National Institute of Health Guide for Care of Laboratory Animals. Beginning on E7.5, pregnant dams received daily subcutaneous injections of 15µL per gram of body weight of either saline or 20% ethanol diluted in saline. Subcutaneous route is used to administer daily injections to pregnant dams because there is no chance of accidentally puncturing growing embryos. Mice were randomly assigned to each group. Prior to injections, mice were anesthetized with isoflurane. On E13.5, the mice also received intraperitoneal injections of 50mg per kg of body weight of bromodeoxyuridine (BrdU) for another study. Thirty minutes after receiving the BrdU injections, mice were euthanized with CO_2_. Embryos were removed via Caesarian section and preserved in 4% paraformaldehyde (PFA) at 4°C. Litter is the n and one embryo from each sex, when applicable, was tested from each litter.

### Sex Genotyping

Tails were removed from each embryo for DNA isolation. To distinguish between males and females, the Jarid1C and Jarid1D genes were amplified (Clapcote & Roder 2005). DNA extraction was performed using the Epoch Life Sciences *GenCatch* Genomic DNA Extraction kit (Epoch 2460050-1/13). We followed an adjusted version of the kit’s protocol. The attached protocol suggests using 200µL of heated water or Tris as an elution buffer, but we used 20µL of heated elution buffer (Buffer AE) from the Qiagen DNeasy Blood & Tissue Kit to maximize DNA yield (Qiagen 69506). Once DNA was isolated, concentrations were measured using a Nanodrop 2000 (Thermo Fisher). We prepared a stock solution of sex primers: 1µL of forward primer (5’-CTGAAGCTTTTGGCTTTGAG-3’), 1µL of reverse primer (5’-CCACTGCCAAATTCTTTGG-3’), and 100µL of sterile H_2_O. For each sample, we prepared this mixture for PCR: 10µL of Taq polymerase (Sigma P0982), 1µL of the premade sex primer stock, 200ng of DNA, and enough sterile H_2_O to fill the volume to 20µL. Samples ran through a PCR program of 35 cycles in the following: denaturing for 20 seconds at 94°C, elongation for 60 seconds at 54°C, and extension for 40 seconds at 72°C. Following PCR, the samples were run on a 2% agarose gel diluted in 0.5X TBE at 100V for approximately 90 minutes.

### Microtome Sectioning

One male and one female embryo from each litter were used in this study. After determining the sex, chosen embryos were sent to AML Laboratories (Jacksonville, FL) and embedded in paraffin in the coronal orientation. Some embryos were also sectioned in-house with a Leica microtome. All embryos were sectioned at 5-micron thickness.

### Nissl Staining

E15.5 brains from Swiss Webster mice were preserved by paraffin wax histology and sectioned along the coronal plane. Every fifth slide from each brain was selected for Nissl staining in order to identify appropriate slides for immunostaining containing brain regions of interest. Wax was removed by a 5 minute bath in Histoclear (Electron Microscopy Sciences #64110). Tissue was then rehydrated by a 5 minute bath in 100% ethanol, followed by 2 minute baths in 95% ethanol, 70% ethanol, and distilled water, in that order. Slides were stained by a 2 minute bath in 0.1% cresyl violet acetate (Electron Microscopy Sciences #26089-01), followed by a 2 minute rinse in distilled water. The tissue was then dehydrated by a series of 2 minute baths in 70% ethanol and 95% percent ethanol, followed by a 5 minute bath in 100% ethanol and a 5 minute bath in Histoclear. Slides were allowed to dry in open air for 1 hour prior to coverslipping with Cytoseal (Electron Microscopy Sciences #18007).

### Immunostaining, Microscopy, and Axon Tracing

E15.5 brains were prepared as previously described. The same tissue rehydration procedure was used. Slides were incubated in citrate antigen retrieval solution (Vector H-3300) in a hot water bath for 20 minutes, followed by a 2 minute incubation in distilled water and a 5 minute incubation in 1X PBS. After these rinses, antigen blocking solution was administered at 200µL per slide, consisting of 10% goat serum and 0.1% Triton (Electron Microscopy Sciences #22145) in 1X PBS. Slides were blocked for 30 minutes, protected by parafilm coverslips to prevent drying. After, coverslips were removed and slides were rinsed for 5 minutes in 1X PBS. Slides were kept in a 1X PBS bath until primary antibody was prepared. Anti-L1 NCAM antibody (AbCam #ab208155) was used at a concentration of 1:250 in 1X PBS, and slides were incubated overnight at 4°C. The next day, the slides were given three 5 minute washes in 1X PBS, then incubated for 30 minutes in 200 μL each of ImmPRESS goat anti-rabbit secondary antibody (Vector #MP-7401). Slides were also covered with parafilm during the incubation. Parafilm was then removed and slides were rinsed for 5 minutes in a 1X PBS bath, then 5 minutes in a distilled water bath. Diaminobenzidine (DAB; Vector #SK-4105) was used to visualize staining. Slides were covered with 200μL each of DAB and incubated for 1 minute before being submerged in a distilled water bath. Slides were then dehydrated as previously described, and allowed to dry before coverslipping. Images were collected using a Nikon Eclipse 80i Compound Microscope at 10X magnification equipped with a Nikon C2-SH digital camera. NIS-Elements AR software was used to capture images. L1-stained axons from each side of section were analyzed using ImageJ with AxonTracer (Patel et al. 2018). Students were not aware of sex or treatment during analysis.

## Acknowledgements

We would like to extend our gratitude to Avery Sicher for her assistance with Axon Tracer and Favero lab members (Rahsaan Sailes, Kevyn Dewees, Deborah Diaz, Erin Mathews, Betty Endeshaw, Hira Khattak, Katherine Renzi, Laura Rothschild, Will Klinger) for their help with all lab procedures, such as brain sectioning, sex genotyping, and coverslipping. Likewise, we would like to thank Dr. Carlita Favero for both lab oversight and the hands-on work with embryo retrieval and mouse C-sectioning. Due to the collaborative nature of this study, we would also like to acknowledge our class peers and lab group members for their work throughout this experiment. Lastly, our gratitude goes out to both the Biology Department and Neuroscience Program for giving us the opportunity to pursue this project.

## Funding

This work was partially supported by NIAAA grant #1R21AA025740-01A1 to C.B.F.

## Notes

### Competing Interest Statement

The authors have declared no competing interest.

## References

Chi, P., Aras, R., Martin, K., Favero, C. (2015). Using Swiss Webster mice to model Fetal Alcohol Spectrum Disorders (FASD): An analysis of multilevel time-to-event data through mixed-effects Cox proportional hazards models. Behav Brain Res 305, 1-7. PMID: 26765502

Clapcote, S. J., Roder, J. C. (2005). Simplex PCR assay for sex determination in mice. BioTechniques 38(5), 702-706. PMID:15945368

De Bellis M.D., Narasimhan A., Thatcher D.L., Keshavan M.S., Soloff P., Clark D.B. (2005). Prefrontal cortex, thalamus, and cerebellar volumes in adolescents and young adults with adolescent-onset alcohol use disorders and comorbid mental disorders. Alcohol Clin Exp Res 29(9), 1590-1600. PMID:16205359

Garel S., López-Bendito G. (2014). Inputs from the thalamocortical system on axon pathfinding mechanisms. Curr Opin Neurobiol 27, 143–150. PMID:24742382

Godin, E. A., O’Leary-Moore, S. K., Khan, A. A., Parnell, S. E., Ament, J. J., Dehart, D. B., Johnson, B. W., Allan Johnson, G., Styner, M. A., & Sulik, K. K. (2010). Magnetic resonance microscopy defines ethanol-induced brain abnormalities in prenatal mice: Effects of acute insult on gestational day 7. Alcoholism: Clinical and Experimental Research. https://doi.org/10.1111/j.1530-0277.2009.01071.x

Granato, A., Santarelli, M., Sbriccoli, A., Minciacchi, D. (1995). Multifaceted alterations of the thalamo-cortico-thalamic loop in adult rats prenatally exposed to ethanol. Anat Embryol 191(1), 11-23. PMID:7717529

Grove, E. A. (2005). Local axon guidance in cerebral cortex and thalamus: Are we there yet? Neuron 48(4), 522-524. PMID:16301165

Harding A., Halliday G., Caine D., Kril J. (2000) Degeneration of anterior thalamic nuclei differentiates alcoholics with amnesia. Brain. 123, 141-154. PMID:10611128

Lebel C., Rasmussen C., Wyper K., Walker L., Andrew G., Yager J., Beaulieu C. (2008) Brain diffusion abnormalities in children with fetal alcohol spectrum disorder. Alcohol Clin Exp Res 32(10), 1732-1740. PMID:18671811

Lindsley, T. A., Shah, S. N., Ruggiero, E. A. (2011) Ethanol alters BDNF-induced Rho GTPase activation in axonal growth cones. Alcohol Clin Exp Res 35(7), 1321-1330. PMID:21676004

Rodriguez C.I., Davies S., Calhoun V., Savage D.D., Hamilton D.A. (2016) Moderate prenatal alcohol exposure alters functional connectivity in the adult rat brain. Alcohol Clin Exp Res 40(10), 2134-2146. PMID:27570053

Patel A., Li Z., Canete P., Strobl H., Dulin J., Kadoya K., Gibbs D., Poplawski G.H.D. (2018) AxonTracer: a novel ImageJ plugin for automated quatification of axon regeneration in spinal cord tissue. BMC Neurosci 19(1), 8. PMID:29523078

Petrelli B., Weinberg J., Hicks G.G. (2018) Effects of prenatal alcohol exposure (PAE): insights into FASD using mouse models of PAE. Biochem Cell Biol 96(2), 131-147. PMID:29370535

Patel, A., Li, Z., Canete, P., Strobl, H., Dulin, J., Kadoya, K., Gibbs, D., & Poplawski, G. H. D. (2018). AxonTracer: A novel ImageJ plugin for automated quantification of axon regeneration in spinal cord tissue. BMC Neuroscience. https://doi.org/10.1186/s12868-018-0409-0

Schambra, U. (2008). Prenatal Mouse Brain Atlas. In Prenatal Mouse Brain Atlas. https://doi.org/10.1007/978-0-387-47093-1

Sepulveda, B., Carcea, I., Zhao, B., Salton, S. R., Benson, D. L. (2011) L1 cell adhesion molecule promotes resistance to alcohol-induced silencing of growth cone responses to guidance cues. Neuroscience 180, 30-40. PMID:21335065

Sokol, R. J., Delaney-Black, V., Nordstrom, B. (2003). Fetal alcohol spectrum disorder. JAMA 290(22), 2996-2999. PMID: 14665662

